# Modulation of Electrostatic Interactions as a Mechanism of Cryptic Adaptation of *Colwellia* to High Hydrostatic Pressure

**DOI:** 10.1101/2024.07.28.605522

**Authors:** George I. Makhatadze

## Abstract

The role of various interactions in determining the pressure adaptation of the proteome in piezophilic organisms remains to be established. It is clear that the adaptation is not limited to one or two proteins, but has a more general evolution of the characteristics of the entire proteome, the so-called cryptic evolution. Using the synergy between bioinformatics, computer simulations, and some experimental evidence, we probed the physico-chemical mechanisms of cryptic evolution of the proteome of psychrophilic strains of model organism, *Colwellia*, to adapt to life at various pressures, from the surface of the Arctic ice to the depth of the Mariana Trench. From the bioinformatics analysis of proteomes of several strains of Colwellia, we have identified the modulation of interactions between charged residues as a possible driver of evolutionary adaptation to high hydrostatic pressure. The computational modeling suggests that these interactions have different roles in modulating the function-stability relationship for different protein families. For several classes of proteins, the modulation of interactions between charges evolved to lead to an increase in stability with pressure, while for others, just the opposite is observed. The latter trend appears to benefit enzyme activity by countering structural rigidification due to the high pressure.

## Introduction

Hydrostatic pressure is an important environmental variable that plays an essential role in biological adaptation for many extremophilic organisms, so-called piezophiles [1, 2]. On Earth, these organisms generally populate the deep ocean floor where hydrostatic pressure can reach 110 MPa (∼1,100 atm) [3, 4]. Single-cell organisms are not the only ones evolved to live under high hydrostatic pressure. The segmented microscopic animal, tardigrade (“water bear”), can survive pressures up to 6,000 atm in the dormant state [5, 6]. Pompeii worms (*Alvinella pompejana*) are species of polychaete worms that live at high pressure and temperature near hydrothermal vents on the ocean floor. More recently, several species of nematode were identified in the deep terrestrial subsurfaces [7]. Bacterial species have been isolated from 1,350 meters inside the Earth’s crust where the temperature reaches 102°C and pressure is estimated to be more than 3,000 atm [8-10]. There are also reports of prokaryotic organisms living at the bottom of oil-well sediments and deep in the Arctic ice [11, 12]. These examples suggest the possibility of life forms on other planets even though the temperature and pressure conditions can be dramatically different from those on the surface of Earth. Such different temperature and pressure conditions are expected to be found under the ice crust on Mars, Jupiter’s moon Europa, and Saturn’s moon Enceladus [13-15].

What are the physico-chemical implications for adaptation to high hydrostatic pressure? There is increasing evidence that there is a difference in how hydrostatic pressure affects the fitness of an organism as opposed to other environmental factors, such as temperature, pH, or salinity [16-18]. To evaluate the effect of one of the factors, the rest of them must be ideally kept constant. The pH and salinity of the ocean remain relatively constant, but the temperature and pressure can vary significantly. However, due to the thermocline, the majority of the organisms that live below 200-300 m meters will experience temperatures ∼4-5°C which are considered psychrophilic conditions [19]. Therefore, the psychrophilic organisms that live at different depths (i.e. with increased piezophilicity) will satisfy the conditions of living at relatively similar temperatures, salinity, and pH and yet have to adapt to different hydrostatic pressures. Marine psychrophilic bacteria from the same genus that inhabit environments with very similar pH, salinity, and temperature, but at different depths, can serve as a model to study the adaption mechanisms to different hydrostatic pressures.

### Selection of Model Organism

The majority of recognized psychropiezophilic strains are found within phylogenetically specific branches of the *Gammaproteobacteria* class. These branches include various notable genera such as *Colwellia, Shewanella, Moritella, Photobacterium*, and *Psychromonas* [17]. Particularly, some of the most extreme psychrophilic and piezophilic species have been identified within the *Colwellia* genus [20]. Members of the *Colwellia* genus are heterotrophic and facultative anaerobic bacteria [21].

One of the first known obligate piezophiles was isolated from the amphipod *Hirondellea gigas* from the Mariana Trench at a depth of 10,476 m [22]. This strain, *Colwellia marinimaniae* sp. MT41, turned out to be psychrophilic that in the lab grows only at pressure above 35 MPa with the optimum growth at 103 MPa [19, 22, 23]. More recently, another *Colwellia marinimaniae* MTCD1, the most piezophilic microbe known to date, was isolated from an amphipod collected from the Mariana Trench at a depth of 10,918 m [24]. This strain displays the growth rate dependence even more extreme than MT41: MTCD1 grows only above 80 MPa up to 140 MPa with an optimum growth pressure of 120 MPa. Another piezophilic and psychrophilic strain of *Colwellia, Colwellia* sp. TT2012 was isolated from the sediments collected at 9,161 m in the Tonga Trench and showed an optimal growth pressure of 84 MPa [20]. A moderate psychropiezophilic strain of *Colwellia*, ATCC-BAA-637, was isolated from the sediments in the Japan Trench at a depth of 6,278 m [20]. It shows growth between 40 and 80 MPa with the optimal growth at 60 MPa. *Colwellia psychrerythraea* GAB14E, a moderate psychropiezophilic strain, was collected in the Great Australian Bight at a depth of 1,472 m [20]. Finally, two piezotolerant *Colwellia* strains, *C. psychrerythraea* 34H, a psychrophile isolated from Arctic Ocean sediments at 305 m and *C. psychrerythraea* ND2E, collected in the Mediterranean Sea from a depth of 495 m [20]. For comparison, with the psychropiezophilic strains listed above, we can use psychrophilic *Colwellia* collected from surface fractions of Arctic seawater *Colwellia* sp. Arc7-635 [25] and *Colwellia polaris* isolated from Arctic sea ice [26]. We thus selected these nine *Colwellia* strains (marine psychrophilic bacteria) as model organisms to elucidate the adaptation to high hydrostatic pressure from the comparative analysis of their proteomes.

### Evidence of Cryptic Evolutionary Adaptation to Hydrostatic Pressure in the *Colwellia* strains

The phylogenetic relationship between nine *Colwellia* strains, based on comparative analysis of the whole genomes (Figure 1A) or the 16S RNA sequences (Figure 1B), shows a clear segregation of strains based on the isolation depth. Since these strains of *Colwellia* have been isolated from the samples collected at different depths but from similar temperature environments, these strains should display the same temperature tolerance profile but different tolerance to pressure as assessed by the growth characteristics discussed above [20]. This allows the use of isolation depth as a proxy for the pressure tolerance of these organisms [16]. The segregation of genomes of *Colwellia* strains observed based on their isolation depth suggests the potential existence of certain physico-chemical characteristics within an organism’s entire proteome. These characteristics might provide an increase in resistance to pressure. Such behavior, as a mechanism of general adaptation to extreme conditions termed **<u>cryptic evolution</u>**, has been discussed in the literature before [27, 28] and several instances where it is observed have been reported [18, 29-31].

**Figure 1.**
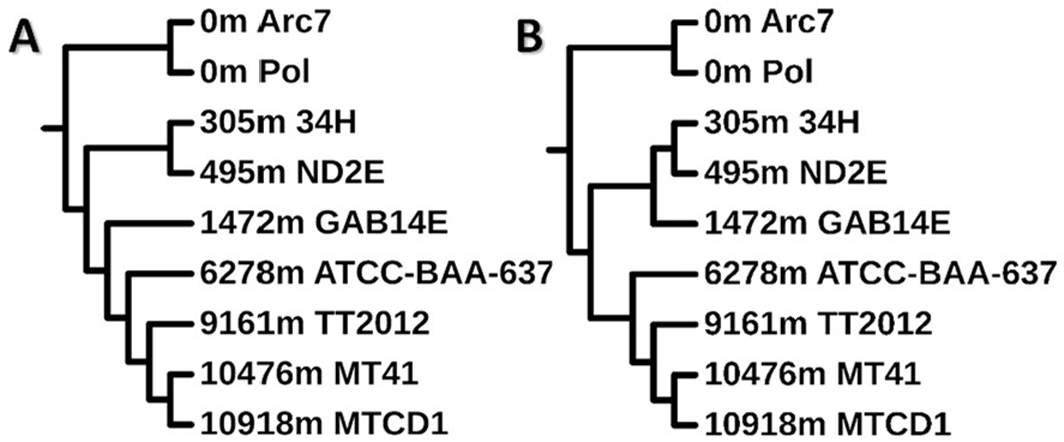
Phylogenetic trees for nine psychrophilic *Colwellia* strains based on the whole genome (A) or 16S RNA (B) analysis [20]. The number ending with the letter “m” indicates isolation depth in meters, followed by the *Colwellia* strain name.

An indication that cryptic evolution might define phylogenetic relationships of the nine analyzed strains of *Colwellia* comes from a general physico-chemical analysis of these proteomes. We computed the isoelectric points, pI, of all proteins in each proteome using sequence information only and pKa values of free amino acids [32], i.e. not accounting for possible shifts in individual pKa values due to interactions that are imposed by the 3D structure. We also computed pI values of all proteins in these proteomes using PROPKA [33, 34] and the 3D structures that we modeled by ESMFold [35]. The results are very consistent (Figure 2A, 2B) and show that the fraction of proteins with acidic pI (defined as pI ≤ 7.6) is lower in most piezophilic strains (MT41 or MTCD1 61±1% computed from sequence and 58±1% from 3D) than in non-piezophilic strains (CP or C-Arc7 70±1% computed from sequence and 66±1% from 3D). Moreover, there appears to be a linear dependence of the fraction of proteins with acidic pI on the isolation depth (i.e. the parameter that we use as a proxy for hydrostatic pressure Figure 2B). In physico-chemical terms, the prevalence of proteins with acidic pI suggests a net excess of acidic residues over basic. It appears that this indeed is the case (Figure 2C): the fraction of acidic residues (D, E) decreases with the isolation depth while the fraction of basic residues (R, K) increases, resulting in an overall decrease in the DE/RK ratio with increase in isolation depth.

**Figure 2.**
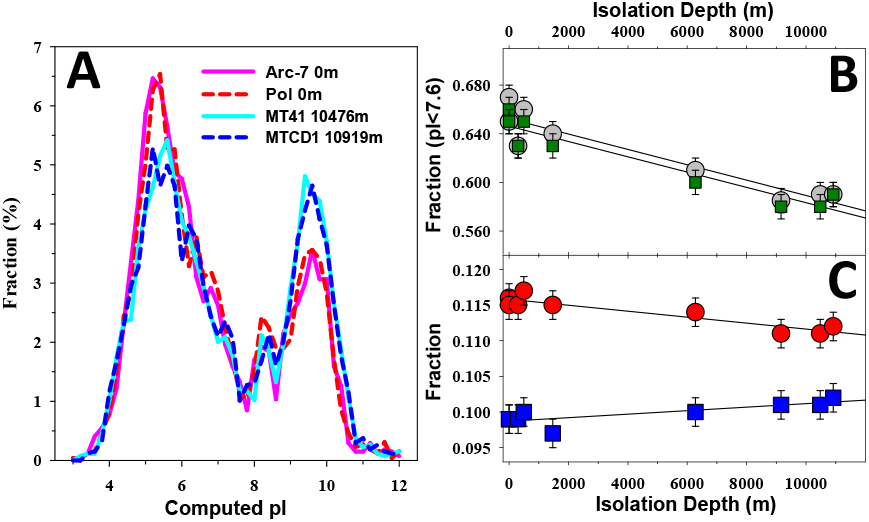
Analysis of the distributions of pI in proteomes of nine Colwellia strains. Panel A. Histograms for pI in non-piezophilic (pink & red) and most piezophilic (blue & cyan) strains. Panel B. Fraction of proteins with pI<7.6 as a function of isolation depth: 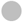 computed directly from the sequences; 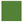 computed from 3D structures modeled using ESMFold. Panel C. changes in fraction of acidic (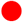 - D & E) and basic (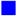 - R & K) residues as a function of isolation depth.

### Manifestation of Cryptic Evolution in Individual Protein Families

To summarize, nine *Colwellia* strains considered here have defined phylogenetic trees, whole genome or 16S RNA-based, that segregate strains according to their isolation depth. Moreover, *the individual proteomes show differences in the fraction of charged residues that also track with the isolation depth*. The next question to ask is do the individual protein families show phylogenetic relationships at the sequence level that also track with the isolation depth? To answer this question we looked at several biophysically tractable protein families: cold shock protein - CspD, ribosomal protein - L30, acylphosphatase - ACP, and dihydrofolate reductase - DHFR. Of these ACP and DHFR are enzymes that possess catalytic activity that is required for organismal survival and were proposed to be present in the last universal common ancestor - LUCA [36-38]. All these four protein families show sequence variation that appears to track with the isolation depth, suggesting that they evolved to adapt to function under increasing pressure. Indeed, sequence-based phylogenetic trees for these proteins show segregation based on the isolation depth (Figure 3). Importantly, their phylogenic trees are similar to the trees obtained using whole genome or 16S RNA for these nine strains of *Colwellia* (see Figure 1).

**Figure 3.**
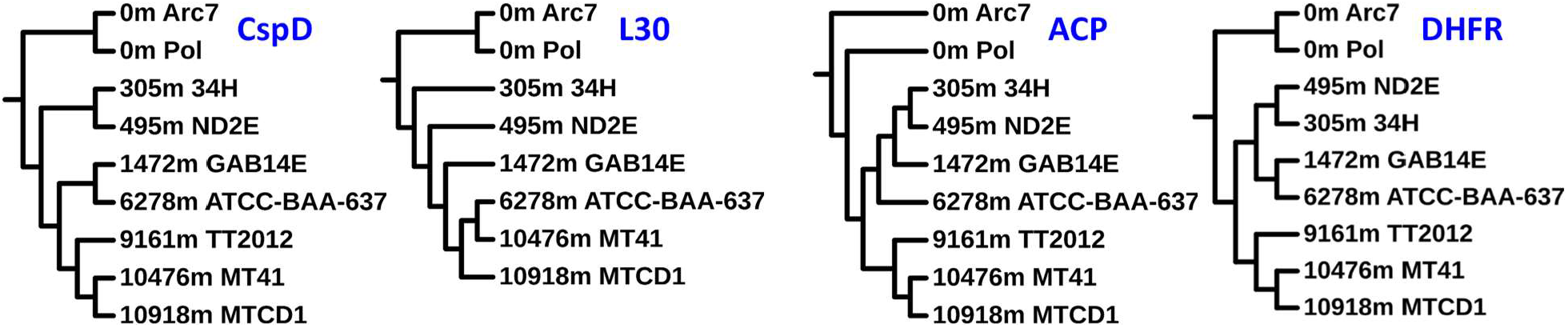
Phylogenetic trees for CspD, L30, ACP, and DHFR based on the sequences from nine (eight in the case of L30) strains of psychrophilic *Colwellia* computed using ClustalO [39].

It is well established that the increase in hydrostatic pressure decreases protein stability and high enough pressure leads to protein unfolding [17, 40]. So one possible mechanism of adaptation will be the increase in the protein stability. Moreover, since we have observed that at the whole proteome level, there are ***changes in the relative fractions of acidic and basic residues***, it is reasonable to hypothesize that ***interactions between ionizable residues***, the so-called charge-charge interactions, might be modulating the properties, such as stability [41-43], of these proteins as a mechanism of pressure adaptation.

We performed the analysis of interactions between charged residues using the protocol developed previously by us and tested for a large number of different proteins including Csp, L30 and ACP (references [41-57]). The energies of charge-charge interactions, ΔG_qq_, are computed using the TKSA algorithm described by us [45, 58]. It is important to emphasize that the results of TKSA calculations have been compared to the results of other continuum electrostatic models such as H++ [59], MCCE [60, 61], UHBD [62], MM_SCP [63], PDLD [64] and found to produce qualitatively similar results (see e.g. [50, 51, 54, 57)]. Comparative analysis of the dependencies of the energy of charge-charge interactions, ΔG_qq_, for each of the four protein families (CspD, L30, ACP, DHFR) from nine *Colwellia* strains shows striking differences and similarities (Figure 4). Dramatic differences are observed in the dependence of ΔG_qq_ on the isolation depth for ACP and DHFR versus CspD and L30. For ACP and DHFR, ΔG_qq_ values increase with the increase in isolation depth (become less negative) suggesting that the energy of charge-charge interactions in these proteins become less stabilizing with the increase in isolation depth. For CspD and L30, ΔG_qq_ values decrease with the increase of isolation depth (becoming more negative), suggesting an increasing contribution to stability from the charge-charge interactions. Implications of these possible mechanisms of *cryptic evolution* led to specific hypotheses by which these two groups of proteins evolved their sequences for functional adaptation to the increase in hydrostatic pressure.

**Figure 4.**
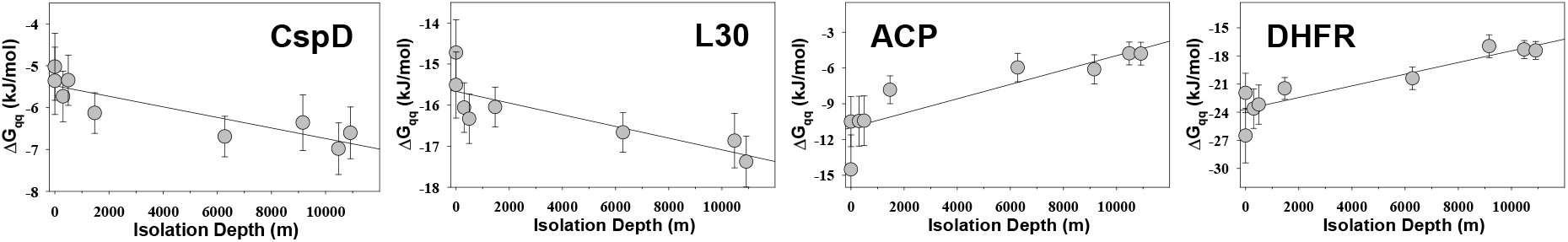
Dependence of energies of charge-charge interactions (ΔG_qq_) on the isolation depth of CspD, L30, ACP and DHFR from nine (eight in the case of L30) strains of psychrophilic *Colwellia*.

### Different Functional Mechanisms of the Piezophilic Adaptation Based on the Cryptic Evolution of Electrostatic Interactions

The contribution of charge-charge interactions to the stability of CspD ***increases*** with an increase in the piezophilicity of the parent strain (Figure 4). As we have shown before for the highly-homologues family of CspB proteins, optimization of charge-charge interactions is a major driving force for an increase in thermostability in this protein family [50, 54]. We have also shown that the increase in pressure stabilizes interactions of CspB proteins with the ssDNA, i.e. the volume change upon CspB:ssDNA binding is positive [65]. Briefly, the following linked equilibrium is modulated by an increase in pressure (knowing that the ligand does not bind to the unfolded state):

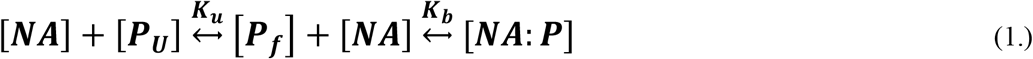

where [P_U_], [P_f_], [NA], and [NA:P] are the free species of unfolded and native protein, DNA, and protein-DNA complex at equilibrium, respectively; K_u_ describes the equilibrium constant for protein stability, K_b_ describes the equilibrium constant of binding. We have shown that the first equilibrium between the unfolded and native states shifts toward the unfolded state with the increase in pressure (i.e. supporting the notion that the unfolded state has a smaller volume than the native state [66, 67]). In the second equilibrium, binding nucleic acid to the protein is shifted toward [NA:P] the increase in pressure suggesting that the binding affinity increases with the increase in pressure. *Thus, the overall bound fraction (i*.*e. [*NA:P*]) as a function of the increase in pressure is largely controlled by the protein stability* [65]. Considering that the DNA binding interface of CspD proteins is almost identical to that of CspB proteins, it is reasonable to assume that to maintain biological function, which is a nucleic acid binding [46-48, 68], the protein needs to be more stable at high pressure [65]. Considering that the correlations for L30 look similar to that for CspD (Figure 4), we postulate that L30 being an RNA binding protein will have the thermodynamic binding signature as CspD, i.e. that the binding equilibria as a function of pressure is controlled by the stability of L30. The number of positions that carry substitutions within the CspD or L30 families is relatively small. Out of 72 residues in CspD, there are 11 positions in which at least one of the nine Colwellia CspD sequences has a substitution. Among these 11 positions, 8 involve substitutions to or from charged residues. Similarly, for the L30 protein family, which is 59 amino acids long, there are 10 positions that can differ in at least one L30 sequence. Of these, 7 are substitutions to or from charged residues. Importantly, all the variable positions are located away from the parts of these two proteins that are involved in the interactions with the nucleic acids (see Figure 5). Thus overall binding equilibrium as a function of pressure is modulated by the stability of CspD or L30: the higher the protein stability is at elevated pressure the higher will be the fraction of the protein:ssDNA complex.

**Figure 5.**
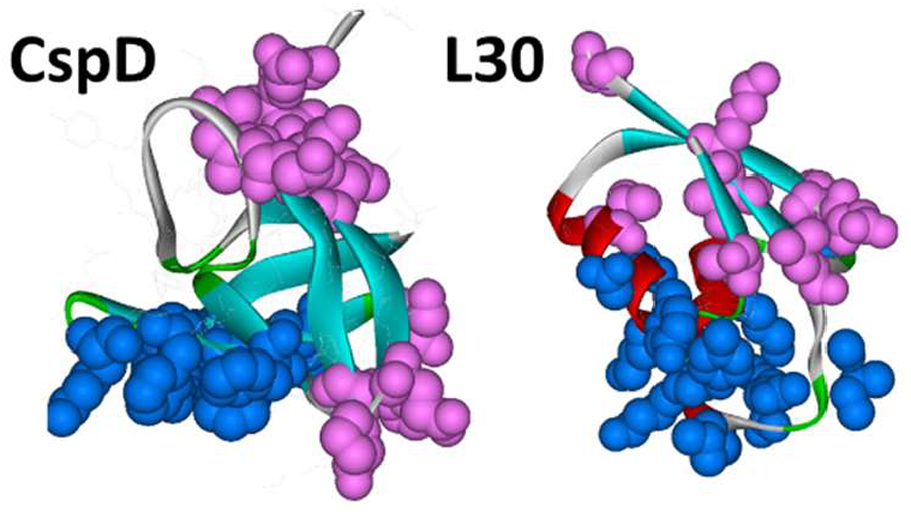
Cartoon structures of CspD and L30 with the positions of substitution sites are shown in pink and residues involved in binding nucleic acids in blue.

The stability of ACP and DHFR from nine *Colwellia* strains isolated at different depths ***decreases*** with an increase in piezophilicity of the parent strain and this decrease in stability is modulated by charge-charge interactions (Figure 4). This decrease in stability is necessary to counter the decrease in the native state dynamics due to the compression of the native state at high pressure. An increase in the hydrostatic pressure induces compression of the native state leading to the decrease in the functional dynamics that is critical for the enzymatic activity. The changes in the native state dynamics are directly related to the enzymatic activity of proteins [69]. To compensate for this decrease in dynamics at higher pressure, there is a decrease in the stability thus restoring the levels of enzymatic activities. This decrease in stability is modulated by the decrease in the strength of charge-charge interactions[68] as can be seen from the progressively less stabilizing values of ΔG_qq_ for proteins from the strains isolated at higher depth for ACP and DHFR (Figure 4). Thus, charge-charge interactions will modulate the recovery of activity of these enzymes at high pressure by providing overall destabilization.

The specific activity of acylphosphatase decreases with the increase in pressure. Figure 6A shows the enzyme concentration normalized initial velocity, V_o_/[E] of hydrolysis of benzoylphosphate at 25°C by human acylphosphatase as a function of pressure from 0 to 2000 bars. It is evident, that the V_o_/[E] decreases with the increase in pressure. This decrease is not caused by enzyme unfolding: Figure 6B shows the effects of hydrostatic pressure on the intrinsic fluorescence of the same enzyme, in the same buffer used for the activity assay, in the presence of 2M glutamate (a cosolvent that leads to an increase in protein stability e.g. [65, 70]) and 6M urea (a cosolvent that leads to the unfolding of a protein). The overlap of fluorescence signal in the absence and presence of 2M glutamate suggests that the enzyme remains folded up to 3,000 bars, i.e. 50% higher pressure than is accessible in the experimental set-up for the activity instruments. This provides a strong indication that the decrease in intrinsic activity upon the increase in pressure is not due to a decrease in protein stability.

**Figure 6.**
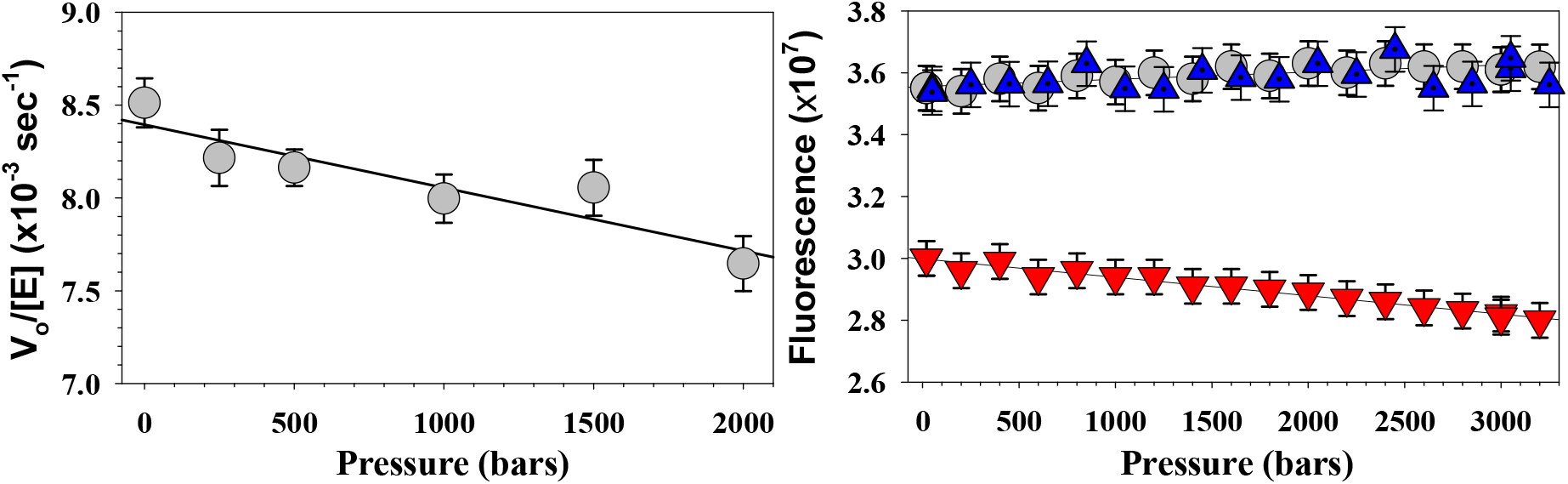
Panel A. Dependence of normalized initial reaction velocity with benzoyl phosphate on pressure for acylphosphatase at 25°C. Panel B. Intrinsic fluorescence intensity as a function of pressure for acylphosphatase in enzyme assay buffer in the absence 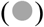 or presence of 2M Glu 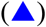or 6M urea 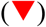. Note the difference in the pressure range in the two panels.

As we discussed above, the number of positions that can carry substitutions within the CspD and L30 families of proteins from *Colwellia*, isolated from different depths, are rather small (<17%), and are somewhat segregated from the functional nucleic acid binding face of the molecule. This is very different for ACP (89 amino acid residues) and DHFR (120 amino acid residues). For these proteins, ∼40% and 47% of positions, respectively, are subject to substitutions. Of those, none are in the proximity (<10Å) of the active site (Figure 7). Moreover, two-thirds of positions are with the substitutions involving charged residues. This observation supports the hypothesis that charge-charge interactions modulate protein stability, and are not directly influencing the enzymatic activity of ACP and DHFR as a function of pressure.

**Figure 7.**
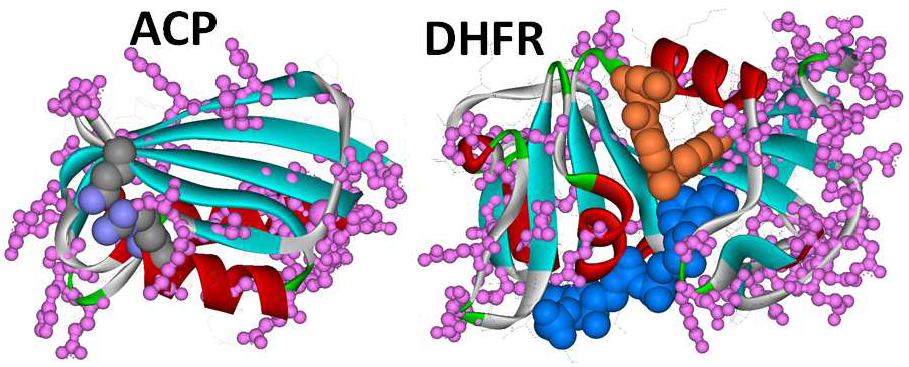
A cartoon representation of the structures of ACP and DHFR with the positions of the substitution site is shown with the pink ball-&stick model. CPK shows the residues in the active site of ACP or the location of the substrates on DHFR.

### Conclusions

The hypothesis of cryptic evolution involving charge-charge interactions outlined here awaits direct experimental validation. The predictions are that the stability of CspD and L30 proteins will correlate directly with the isolation depth, i.e. proteins from piezophilic *Colwellia* strains will be more stable than the ones from the *Colwellia* found at ambient pressures. At the same time, the nucleic acid binding affinity for all proteins from these strains should be largely independent of pressure. The predicted outcomes for the stability of ACP and DHFR proteins from these *Colwellia* strains will be an anti-correlated dependence of stability on the isolation depth. In terms of enzymatic activities, it is expected that all enzymes will have rather similar activities at the corresponding optimal growth pressure.

## Material and Methods

Three-dimensional structures of all proteins in the studied *Colwellia* proteomes were modeled using ESMFold [35] LLM. In addition, sequences corresponding to the selected set of protein families (ACP, DHFR, CspD, L30) were also modeled using AlphaFold2 [71].

The isoelectric point of each protein in a given proteome was computed in two different ways:

1. Directly from the amino acid composition, assuming the non-interacting charges as implemented in IPC2.0 [72].
2. From the modeled 3D structures of proteins using PROPKA3 [33]. In this case, the effects of interactions between charges on the individual pKa and its contributions to the pH dependence of the ionization state (and thus the computed pI) were accounted.

To facilitate comparison between the two methods, the pKa values for IPC were set to unperturbed values used in PROPKA3 (i.e. Asp – 3.0; Glu – 4.5; His – 6.5; Cys – 9.0; Tyr – 10.0; Lys – 10.5; Arg – 12.5; C-term – 3.2; N-term – 8.0).

Analysis of charge-charge interactions according to the TKSA formalism were performed as described by us before [41, 45, 73]. The starting structures for the analysis were obtained using AlphaFold2 (see above).

The activity of ACP as a function of pressure using a High-Pressure Stopped Flow Spectrophotometer/Fluorimeter system (TgK Scientific Limited, Bradford-on-Avon, UK) instrument. The activity was measured at 25°C in 100 mM NaAcetate, pH 5.5 buffer using synthetic substrate benzoyl phosphate (BP). Hydrolysis of BP was followed spectrophotometrically, by measuring the decrease in absorbance at 283 nm upon enzymatic hydrolysis of benzoyl phosphate to benzoic acid [74, 75]. Benzoyl phosphate was synthesized as described in [56, 75].

## Author Contributions

GIM performed all the analysis and wrote the paper.

## Data Availability Statement

All data required for the conclusions made here are contained in the article. Any other data is available upon request from the author.

